# An extracellular protein regulates patched-related/DAF-6-mediated sensory compartment formation in *C. elegans*

**DOI:** 10.1101/2020.05.29.123182

**Authors:** Hui Hong, Huicheng Chen, Yuxia Zhang, Zhimao Wu, Yingying Zhang, Yingyi Zhang, Zeng Hu, Jian Zhang, Kun Lin, Jinghua Hu, Qing Wei

**Affiliations:** CAS Key Laboratory of Insect Developmental and Evolutionary Biology, CAS Center for Excellence in Molecular Plant Sciences, Institute of Plant Physiology and Ecology, Chinese Academy of Sciences, Shanghai 200032, China; Department of Thoracic Surgery, Fudan University Shanghai Cancer Center, Shanghai 200032, China; Department of Oncology, Shanghai Medical College, Fudan University Shanghai Cancer Center, Shanghai 200032, China; University of Chinese Academy of Sciences, Beijing 100039, China; Laboratory for Reproductive Health, Institute of Biomedicine and Biotechnology, Shenzhen Institutes of Advanced Technology, Chinese Academy of Sciences (CAS), Shenzhen 518055, China; Department of Biochemistry and Molecular Biology, Mayo Clinic, Rochester, Minnesota 55905, USA; Department of Cancer Biology, UT MD Anderson Cancer Center, Houston, TX 77230, USA

**Author notes:** These authors contributed equally to this work (Hui Hong, Huicheng Chen).

## Abstract

Coordination of neurite extension with surrounding glia development is critical for neuronal function, but the underlying molecular mechanisms remain poorly understood. Through a genome-wide mutagenesis screen in *C. elegans*, we identified *dyf-4* and *daf-6* as two mutants sharing similar defects in dendrite extension. DAF-6 encodes a glia-specific patched-related membrane protein that plays vital roles in glial morphogenesis. We cloned *dyf-4* and found that DYF-4 encodes a glia-secreted extracellular protein. Intriguingly, DYF-4 colocalizes with DAF-6 along the glial channel. Further investigations revealed that DYF-4 directly interacts with DAF-6 and regulates its proper membrane localization. Notably, reported glial suppressors of *daf-6* could also restore dendrite elongation and ciliogenesis in both *dyf-4* and *daf-6* mutants. Collectively, our data suggest that secreted DYF-4 likely acts as a novel ligand/regulator for the patched-related receptor DAF-6 which promotes the proper formation of the glial channel and indirectly affects neurite extension and ciliogenesis.

## Introduction

The sensory organ uses its ending to receive environmental stimuli. In many cases, the ciliated neuronal receptive endings and surrounding glial cells form compartmentalized sensory endings, as observed in *C. elegans*, *Drosophila* and the olfactory epithelium of mammals (Burkitt G. H, 1993; Kernan, 2007; Ross M. H, 1995; Shaham, 2010). The proper formation of the sensory compartment requires interplay between glial cells and the ciliary endings of sensory neurons. The integrity of the glial compartment and the development of sensory neurons impact each other in a reciprocal manner (Bacaj et al., 2008; Heiman and Shaham, 2009; Singhal and Shaham, 2017). However, little is known about the underlying mechanism.

*C. elegans* uses its sensory organs (amphid and phasmid) to sense environmental cues. The amphid sensory organ of the head consists of 12 sensory neurons, whose ciliated endings are wrapped by the ends of a sheath glia cell and a socket glia cell (Inglis et al., 2007). The cell bodies of sensory neurons are located in the pharyngeal bulb, with the dendrites extending to the anterior end of the nose and terminating with cilia. Both the sheath and socket glial cells extend in parallel along the dendrites of sensory neurons and form discrete tubular channels surrounding amphid cilia. Specifically, the proximal portion of cilia is surrounded by the membrane of the sheath cell, while the distal portion of cilia extends through a pore formed by the membrane of the socket cell. The structure of the phasmid of the tail is similar to that of the amphid, except that it contains only 2 sensory neurons.

The proper formation of the sensory compartment in *C. elegans* includes the compartmentation of glial endings, neuron dendritic anchoring and cilia biogenesis. DAF-6, a patched-related membrane protein, is a key regulator of the compartmentation of the glial endings in *C. elegans* (Perens and Shaham, 2005). DAF-6 functions in the very early stage to restrict the expansion of the glial compartment (Oikonomou et al., 2011), and its activity can be antagonized by several suppressors, including the Nemo-like kinase LIT-1, the actin regulator WSP-1, some retromer components (SNX-1, SNX-3, VPS-29) and the Ig/FNIII protein IGDB-2 (Oikonomou et al., 2012; Oikonomou et al., 2011; Wang et al., 2017). DAF-6 and its suppressors may regulate vesicle dynamics in glial cells in a cell-autonomous manner. In *C. elegans*, dendrite extension in the amphid and phasmid occurs via a special “retrograde extension” process (Heiman and Shaham, 2009; Schouteden et al., 2015). Specifically, dendritic tips are first anchored at the site where the sensory complex will be assembled, and the cell bodies then grow backward to elongate the dendrites. It has been reported that the tectorin-related proteins DEX-1 and DYF-7 form an anchorage in the extracellular matrix and play essential roles in dendritic anchoring (Heiman and Shaham, 2009). The transition zone at the ciliary base is also involved in dendritic anchoring in *C. elegans* (Schouteden et al., 2015). Combined mutations of individual genes from the Meckel-Gruber Syndrome (MKS) module and Nephronophthisis (NPHP) module result in severe defects in dendrite elongation (Li et al., 2016; Schouteden et al., 2015; Warburton-Pitt et al., 2012; Williams et al., 2011; Williams et al., 2010). Notably, proper dendritic anchoring and cilia biogenesis are also required for glial compartment formation. Sheath cells fail to undergo extension in *dyf-7* mutants (Heiman and Shaham, 2009). DAF-6 and its suppressors are mislocalized in ciliogenesis mutants (Oikonomou et al., 2011; Perens and Shaham, 2005). However, little is known about the identity and source of the extracellular signals that coordinate the glial channel formation and dendritic morphogenesis.

In a whole-genome forward genetic screen, we identified a novel gene, *dyf-4*, whose mutants recapitulate the phenotypes of *daf-6*, exhibiting truncated dendrites and defective ciliogenesis in both amphid and phasmid neurons. We cloned DYF-4 and found that it encodes a glia-secreted protein that is enriched in the luminal space between glial and ciliary membranes. DYF-4 is indispensable for the surface localization of DAF-6 in the glial compartment. Deletion of *dyf-4* results in abnormal glial compartment formation, which indirectly leads to defects in dendrite elongation and ciliogenesis. Interestingly, the dendritic and ciliary defects of *dyf-4* could be restored by reported *daf-6* suppressors (*wsp-1, lit-1* and *igdb-2*). Our results reveal that DYF-4 and DAF-6 function in the same pathway, likely as a ligand-receptor pair, to regulate the proper formation of the sensory compartment.

## Results

### *dyf-4* encodes *C13C12.2* and is required for dendrite extension in *C. elegans*

In *C. elegans*, amphid cilia and phasmid cilia are wrapped by sheath and socket cells and extend to the external environment. Defects in environmental exposure or ciliogenesis result in dye-filling defects (*dyf*) (Hedgecock et al., 1985). In a forward genetic screen searching for *dyf* mutants, we isolated two mutants, *jhu431* and *jhu500*, sharing identical phenotypes, with extremely short phasmid dendrites and slightly truncated amphid dendrites (Fig. 1A-F; Fig. 2A-F). The shorter dendrites suggest that *dyf* are mainly caused by defects in dendrite elongation in these two mutants. *jhu431* was mapped to *c13c12.2*, a gene of unknown function. C13C12.2 encodes a 383 amino acid protein that contains a signal peptide at its N-terminus and a PLAC (protease and lacunin) domain at its C-terminus (Fig. 1A; Fig. 1-figure supplement 1A and B). The PLAC domain is a six-cysteine region of approximately 40 amino acids that is usually present at the C-terminus of various extracellular matrix proteins (Ahram et al., 2009; James B. Nardi and Robertson, 1999; Somerville et al., 2004). *jhu431* carries a missense mutation affecting one of the conserved cysteines (C366Y) in the PLAC domain (Fig. 1A). A complementation assay indicated that *jhu431* failed to complement *dyf-4(m158)*, a mutant isolated 20 years ago that has not yet been cloned. *dyf-4 (m158)* shows the same phenotype as *jhu431*, including very short phasmid dendrites and slightly shorter amphid dendrites (Fig. 1B-F). Genomic sequencing showed that *dyf-4(m158)* possesses a nonsense mutation (Q166stop) in the middle region of C13C12.2. The elongation defects of the dendrites observed in *dyf-4(m158)* and *jhu431* were fully rescued by the transgenic expression of the wild-type (WT) *dyf-4* gene under the control of its own promoter but were not rescued by the C13C12.3^C366Y^ mutant form (Fig. 1B-F), indicating that C13C12.2 is responsible for these defects and that C366 is critical for its function. Therefore, we refer to C13C12.2 as DYF-4 hereafter.

**Figure 1.**
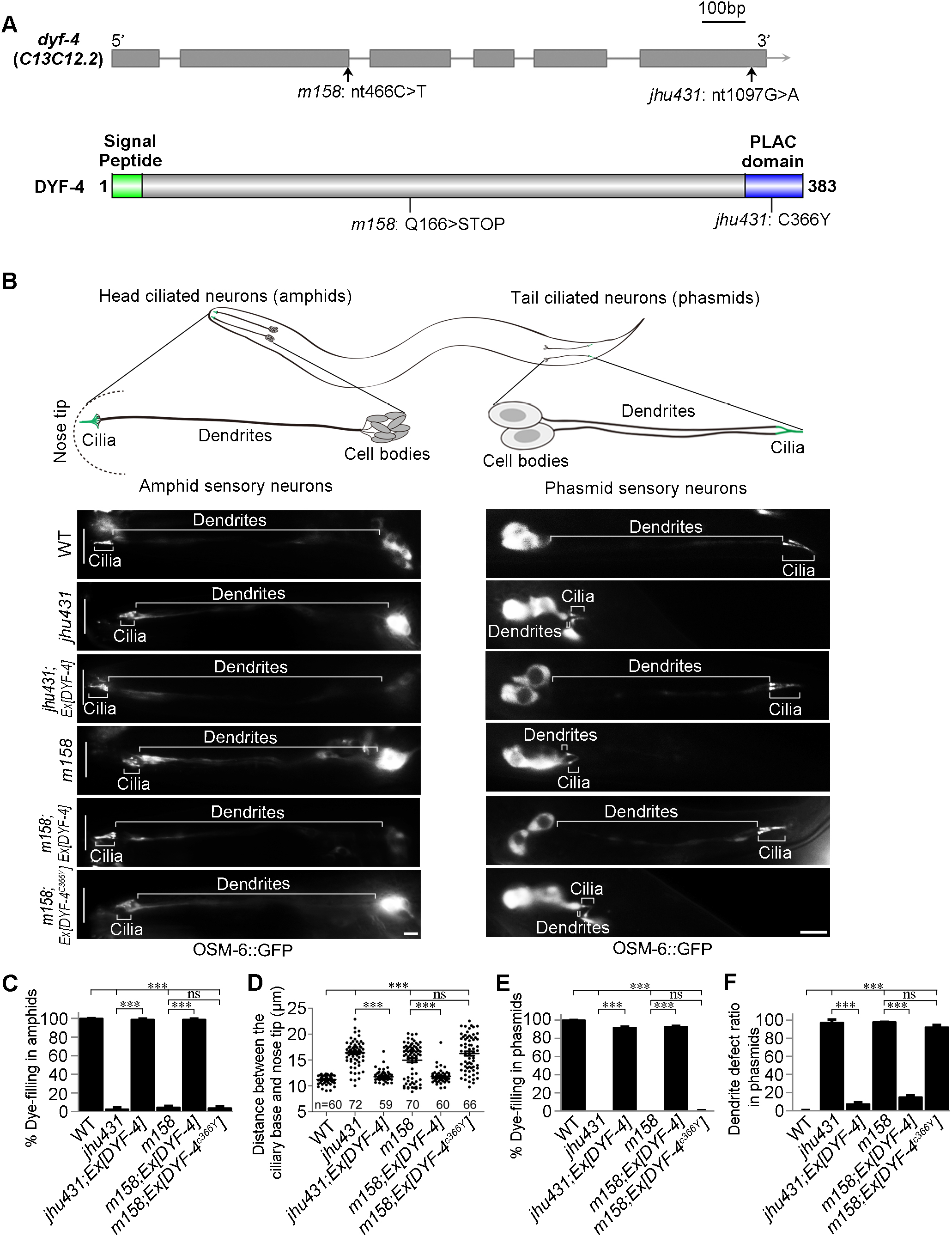
*dyf-4* encodes *C13C12.2* and is required for dendrite extension in *C. elegans*. A. Schematic diagrams of the genomic and protein structures of *dyf-4* (*c13c12.2)*. DYF-4 has a predicted signal peptide at its N-terminus and a PLAC domain at its C-terminus. *dyf-4* (*jhu431)* carries a point mutation (c.1097G>A) that leads to a missense mutation in one of the conserved cysteines (C366Y) in the PLAC domain. *dyf-4*(*m158)* possesses a point mutation (c.466C>T) that results in a nonsense mutation (Q166stop) in the middle region of the DYF-4 protein. As *dyf-4*(*jhu431)* and *dyf-4*(*m158)* exhibit the same mutant phenotypes, we used the *dyf-4*(*m158)* mutant in subsequent experiments if not otherwise specified. B. Upper panels: schematic representation of *C. elegans* amphid and phasmid structures. Lower panels: Fluorescence micrographs of sensory neurons in the amphid and phasmid in WT, *dyf-4*(*jhu431), jhu431*; *Ex[DYF-4]*, *dyf-4*(*m158*), *m158*; *Ex[DYF-4]* and *m158*; *Ex[DYF-4^C366Y^]* worms. The IFT-B component OSM-6::GFP was used to label cell bodies, dendrites and cilia. Scale bars: 5 μm. C. Statistics of the dye-filling percentages in the amphids of the indicated worm lines. D. Statistics of the distance between basal bodies and the nose tip at the head of the indicated worm lines. E. Statistics of the dye-filling percentages in the phasmids of the indicated worm lines. F. Statistics of the phasmid dendrite defect ratio in the indicated worm lines. Data are presented as the mean ± SEM from three independent experiments (n ≥ 80 per experiment). ***P < 0.001 (Student’s t-test).

**Figure 2.**
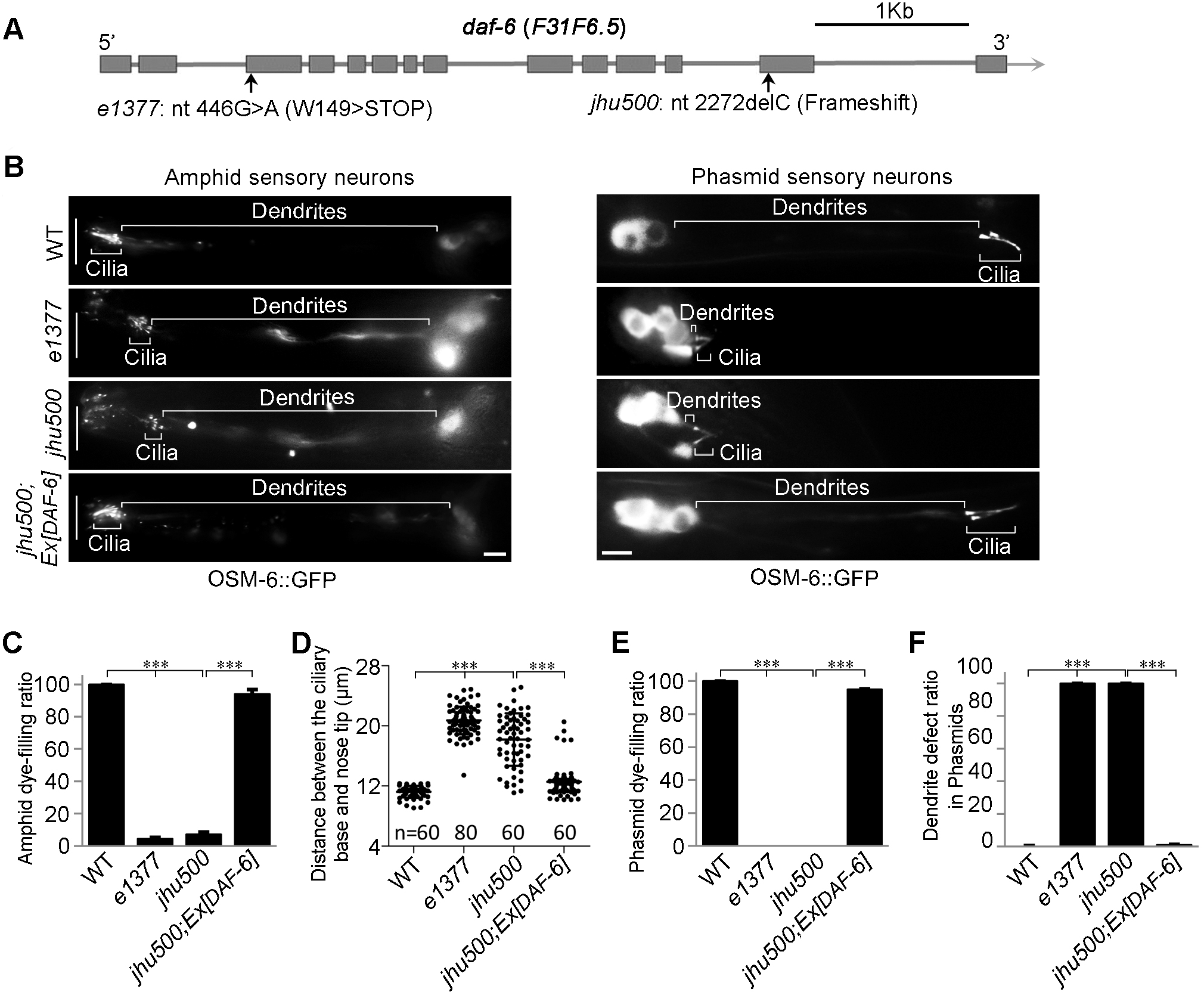
The *daf-6* mutant mimics the phenotype of the *dyf-4* mutant. A. Schematic diagram of *daf-6* alleles. *daf-6(e1377)* was previously reported. *daf-6(jhu500)* has a 1-bp deletion (c.2272delC) at its C-terminus, leading to a frameshift. B. Fluorescence micrographs of sensory neurons in the amphids and phasmids of WT, *daf-6(e1377)*, *daf-6(jhu500)* and *jhu500*; *Ex[DAF-6]* worms expressing OSM-6::GFP. Scale bars: 5 μm. C. Statistics of the dye-filling percentages in the amphids of the indicated worm lines. D. Statistics of the distance between basal bodies and the nose tip at the head in the indicated worm lines. E. Statistics of the dye-filling percentages in the phasmids of the indicated worm lines. F. Statistics of the phasmid dendrite defect ratio in the indicated worm lines. Data are presented as the mean ± SEM from three independent experiments (n ≥ 80 per experiment). ***P < 0.001 (Student’s t-test).

### *daf-6* mimics the phenotype of *dyf-4*

Through a SNP mapping strategy and whole-genome sequencing, the *jhu500* allele was mapped to *daf-6* as a 1-bp deletion (c.2272delC) that leads to a frameshift and loss of the C-terminal region (Fig. 2A). A null allele of *daf-6 (e1377)* showed the same phenotype as *jhu500* and failed to complement *jhu500*. The introduction of the WT *daf-6* cDNA fully rescued the dendrite extension defects of *jhu500*, revealing that *jhu500* is a new allele of *daf-6* (Fig. 2B-F).

DAF-6 is a glial protein similar to the hedgehog membrane receptor patched, which regulates glial compartment morphogenesis in *C. elegans* (Oikonomou et al., 2011; Perens and Shaham, 2005). Our results indicate that DAF-6 is also involved in the dendrite elongation of sensory neurons. Dendrite extension in *C. elegans* is mediated by “retrograde extension”, with dendritic tips first being anchored to the extracellular matrix at the destination, after which the cell bodies travel backward, trailing the elongating dendrites behind them (Heiman and Shaham, 2009). Since the localization of neuron cell bodies is normal in both *daf-6* (*jhu500*) and *dyf-4* (*jhu431*), their short dendrites are caused by defects in dendritic tip anchoring. We speculate that the abnormal glial compartment formation observed in *daf-6* damages the extracellular matrix organization, which indirectly perturbs the anchoring of dendrites. Since the dendritic tips in a sensory compartment are collectively anchored at the destination, the more neurons there are, the stronger the attachment force is (Fan et al., 2019; Schouteden et al., 2015). Therefore, dendrites in amphids (containing 12 neurons) are more resistant to defects than dendrites in phasmids (containing only 2 neurons).

### DYF-4 is a glia-secreted protein that colocalizes with DAF-6 in the sensory compartment

Due to the identical mutant phenotypes shared by *dyf-4* and *daf-6*, we speculate that DYF-4 and DAF-6 function in the same pathway. Hence, we generated a transcriptional reporter for *dyf-4* (*Pdyf-4::*GFP) to determine where *dyf-4* is expressed. We found that *Pdyf-4::*GFP was expressed exclusively in sheath and socket glial cells in both the amphid and phasmid (Fig. 3-figure supplement 1A), suggesting that DYF-4, similar to DAF-6, functions in glial cells rather than sensory neurons to regulate dendrite elongation. To further confirm that glial cells are the functional sites of dyf-4, we expressed *dyf-4* cDNA driven by the *daf-6* promoter or the neuron-specific *dyf-7* promoter. As expected, only the *daf-6* promoter-driven expression of DYF-4 rescued the *dyf-4* mutant phenotype (Fig. 3-figure supplement 1 B).

Next, we examined the subcellular localization of DYF-4 using a GFP-tagged DYF-4 fusion protein. Unfortunately, the DYF-4::GFP signal was too weak to be observed directly. We then employed an indirect immunofluorescence method in which an anti-GFP antibody was used to enhance the DYF-4 signal. Interestingly, DYF-4::GFP was specifically enriched in the sensory compartment region in both amphids and phasmids, primarily surrounding the distal segment of the cilia (Fig. 3A-D). DAF-6 showed a similar localization pattern, with the majority of the signal covering the distal segment of cilia (Fig. 3B-D). The fact that DYF-4 possesses an N-terminal secretion signal peptide and a C-terminal matrix-related PLAC domain led us to propose that DYF-4 is expressed in glial cells and secreted into the extracellular matrix of the sensory compartment. Consistent with this hypothesis, deletion of the signal peptide resulted in the mislocalization of DYF-4 within glial cells (Fig. 3-figure supplement 1C). Notably, the DYF-4^C366Y^ mutation also led to disrupted secretion and accumulation inside glial cells, suggesting that the matrix-related PLAC domain might also regulate the DYF-4 secretion process (Fig. 3-figure supplement 1C).

**Figure 3.**
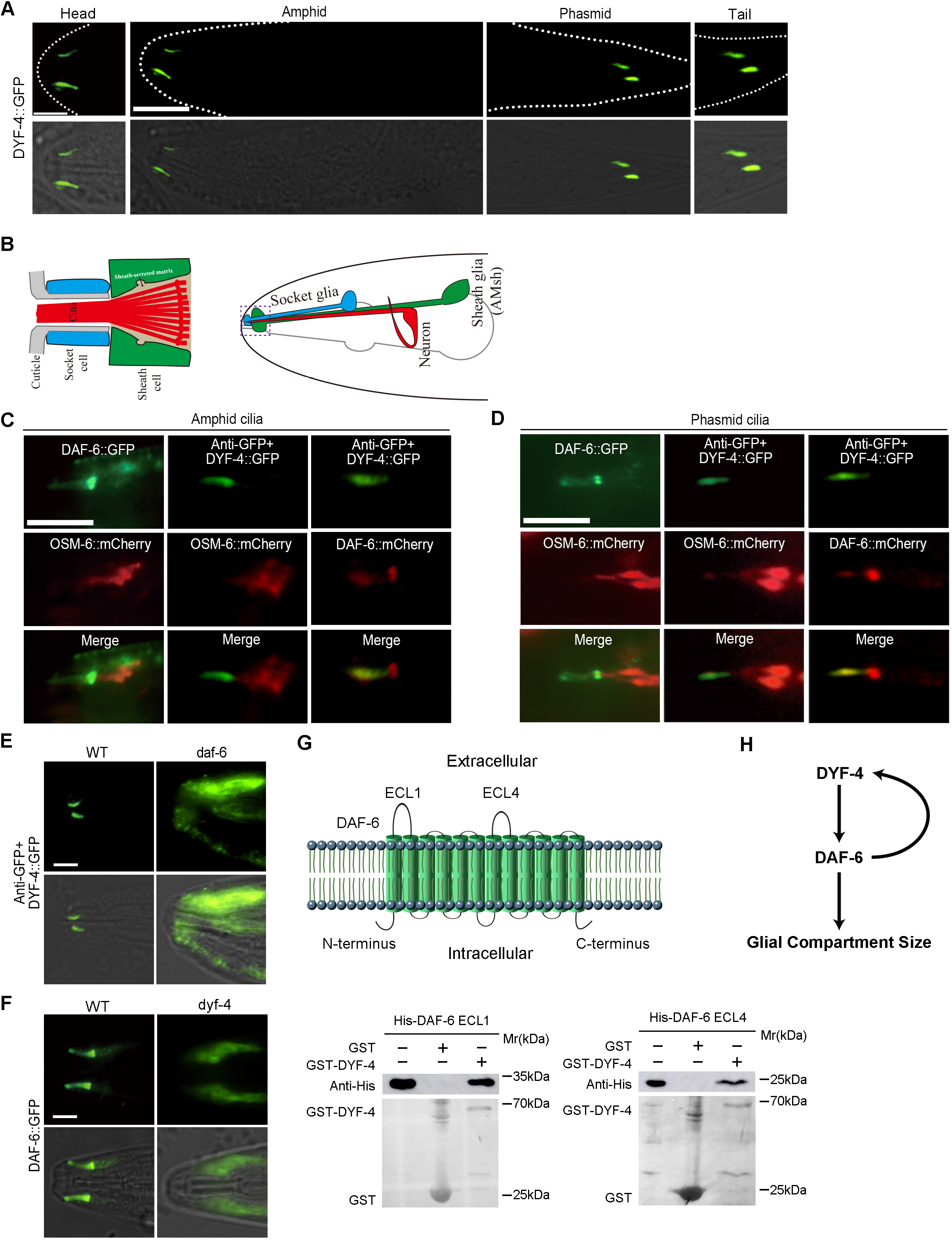
DYF-4 interacts and colocalizes with DAF-6 and is required for DAF-6 localization. A. Representative images of DYF-4::GFP localization in the amphids and phasmids of the WT. DYF-4::GFP localizes to the sensory compartment region. Scale bar: 10 μm. B. Illustration of the *C. elegans* amphid. Right: Each amphid consists of 12 sensory neurons, a socket glial cell and a sheath glial cell. Left: Detail of the sensory compartment in the amphid. C. Colocalization of DYF-4 and DAF-6 along the distal glial channel in the amphid. D. Colocalization of DYF-4 and DAF-6 along the distal glial channel in the phasmid. E. Subcellular localization of DYF-4::GFP in WT and *daf-6* worms. DYF-4 is mislocalized in *daf-6* mutants. F. Subcellular localization of DAF-6::GFP in WT and *dyf-4* worms. DYF-4 is required for the glial channel localization of DAF-6. G. DYF-4 interacts with DAF-6. Upper panel: Schematic representation of DAF-6 transmembrane protein. Lower panel: DYF-4 interacts with both DAF-6 ECL1 and DAF-6 ECl4 in the GST pull-down assay. H. Schematic representation of the relationship between DAF-6 and DYF-4. Scale bars: 5 μm.

### DYF-4 is required for the proper localization of DAF-6 along the surface of the glial channel

Next, we investigated the interdependence of the subcellular localization of the DYF-4 and DAF-6 proteins. Compared to WT, the glial channel localization of DYF-4 was lost, and the protein accumulated inside glial cells in *daf-6* mutants (Fig. 3E), suggesting that DAF-6 is required for DYF-4 secretion. Intriguingly, the surface localization of DAF-6::GFP was also lost in *dyf-4* mutants (Fig. 3F). These data suggest that DYF-4 and DAF-6 depend on each other for transport to and deposition at the cell surface, likely via the same route.

The mislocalization of DAF-6 suggests that glial compartment formation is compromised in *dyf-4* mutants. The cilia of *daf-6* mutants tend to curve laterally due to abnormal glial compartments (Perens and Shaham, 2005). Interestingly, curved cilia were also frequently observed in *dyf-4* amphids (Fig. 3-figure supplement 2A). To further confirm that the glial compartment shows defects in *dyf-4* mutants, we examined two glial channel markers, WSP-1 and ARX-2. Both WSP-1 and ARX-2 localize along the distal portion of the glial compartment channels (formed by socked glia) in *C. elegans* (Oikonomou et al., 2011; Zhu et al., 2017). In *dyf-4* mutants, WSP-1 and ARX-2 could be detected in the sensory compartment but their signal length was significantly shortened (Fig. 3-figure supplement 2B-G), suggesting that the targeting of WSP-1 and ARX-2 to the surface of the glial channel was not affected, but the morphology of glial channels was altered. Considering these findings together, we reasoned that DYF-4 is specifically required for the membrane localization of DAF-6, which then regulates the proper structure/properties of the channel surface.

The similar function, localization, and trafficking defects led us to ask whether DYF-4 physically interacts with DAF-6. DAF-6 is a protein with 12 transmembrane domains, and both its N- and C-termini are predicted to reside in the cytoplasm. On the extracellular side, there are two longer extracellular loops (ECLs), ECL1 and ECL4. We found that DYF-4 could interact directly with both ECL1 and ECL4 (Fig. 3G), supporting the existence of an intriguing mechanism in which a complex formed by a patch-like receptor and a matrix protein is vital for the proper function of glial cells (Fig. 3H).

### *daf-6* suppressors effectively restore *dyf-4* mutant phenotypes

Previous studies demonstrated that the Nemo-like kinase LIT-1, the actin regulator WSP-1, some retromer components and the Ig/FNIII protein IGDB-2 antagonize DAF-6 to promote glial compartment formation. The deletion of these genes also partially restores dye filling defects in the amphids of *daf-6* mutants (Oikonomou et al., 2012; Oikonomou et al., 2011; Wang et al., 2017). We reasoned that if DYF-4 and DAF-6 function together to regulate glial function, *daf-6* suppressors should rescue *dyf-4* phenotypes. As expected, *wsp-1, lit-1* or *igdb-2* mutants all restored the dye-filling defects of *dyf-4* mutants, as they did in *daf-6* mutants (Fig. 4A).

**Figure 4.**
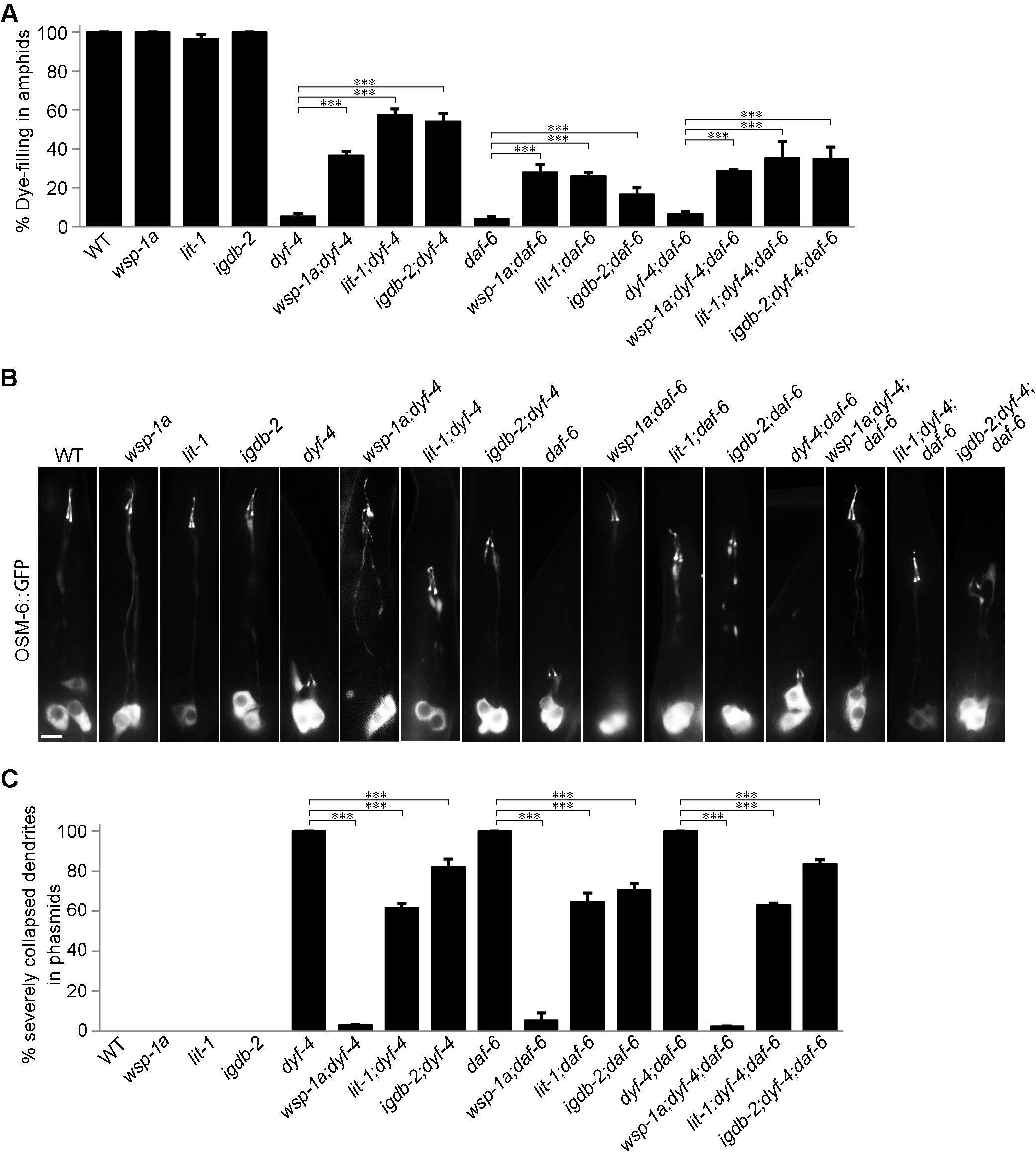
Suppressors of *daf-6* inhibit defects in dendrite extension in both *dyf-4* and *daf-6* mutants. A. Statistics of the dye-filling percentages in the amphid neurons of the indicated worm lines. Suppressors of *daf-6, wsp-1a (gm324)*, *lit-1 (or131)* and *igdb-2* inhibit the defects in dendrite elongation in *dyf-4* and *daf-6* mutants. B. Fluorescence micrographs of phasmid dendrites in the indicated genetic background. Scale bar: 5 μm. C. The percentages of severely collapsed dendrites of the phasmid in the indicated worm lines. Data are presented as the mean ± SEM from three independent experiments (n ≥ 80 per experiment). ***P≤0.001 (Student’s t-test).

To confirm that the restoration of dye filling resulted from the rescue of the shortened dendrites in *dyf-4* and *daf-6* mutants, we focused on phasmids in which dendrites were severely shortened. In *dyf-4* or *daf-6* mutants, all the phasmid neuron dendrites are collapsed into the cell bodies. In contrast, in *dyf-4;wsp-1a* and *daf-6;wsp-1a* double mutants, only 3.19% and 5.55% of phasmid dendrites, respectively, were severely collapsed (Fig. 4B and C). Consistent with these findings, the deletion of *lit-1*(*or131*) or *igdb-2*(*vc21022*) partially inhibited the dendrite elongation defects of *dyf-4* or *daf-6* single mutants (Fig. 4B and C). Notably, the rescue efficiency of *wsp-1* was much higher than that of *lit-1* or *igdb-2*, suggesting that the actin cytoskeleton is the major downstream functional site of DYF-4-DAF-6-regulated glial function.

### The ciliogenesis defects of *dyf-4* and *daf-6* mutants can be rescued by suppressors

In addition to the phenotype of shorted dendrites, ciliogenesis is defective in *dyf-4* and *daf-6* mutants. The cilia are significantly shorter and usually lack distal segments in both types of mutants (Fig. 5A). The length of cilia is approximately 7 μm on average in WT, while it is only approximately 3 μm in both mutants (Fig. 5B). The intraflagellar transport (IFT) machinery is essential for cilia biogenesis (Rosenbaum and Witman, 2002). The examination of various IFT components, including the IFT-B components OSM-6 (the homolog of IFT52) and OSM-5 (the homolog of IFT88), the IFT-A component CHE-11 (the homolog of IFT14) and the BBSome component BBS-7, showed that the ciliary fluorescence levels of all these IFT components were also dramatically decreased, suggesting that the ciliary entry of IFT was compromised in *dyf-4* and *daf-6* mutants (Fig. 5-figure supplement 1). Notably, the observed ciliogenesis defects could be fully rescued by the transgenic expression of the wild-type *dyf-4* or *daf-6* genomic sequence (Fig. 5A and B). How glial dysfunction indirectly impacts IFT behavior and ciliogenesis in sensory neurons remains an open question.

**Figure 5.**
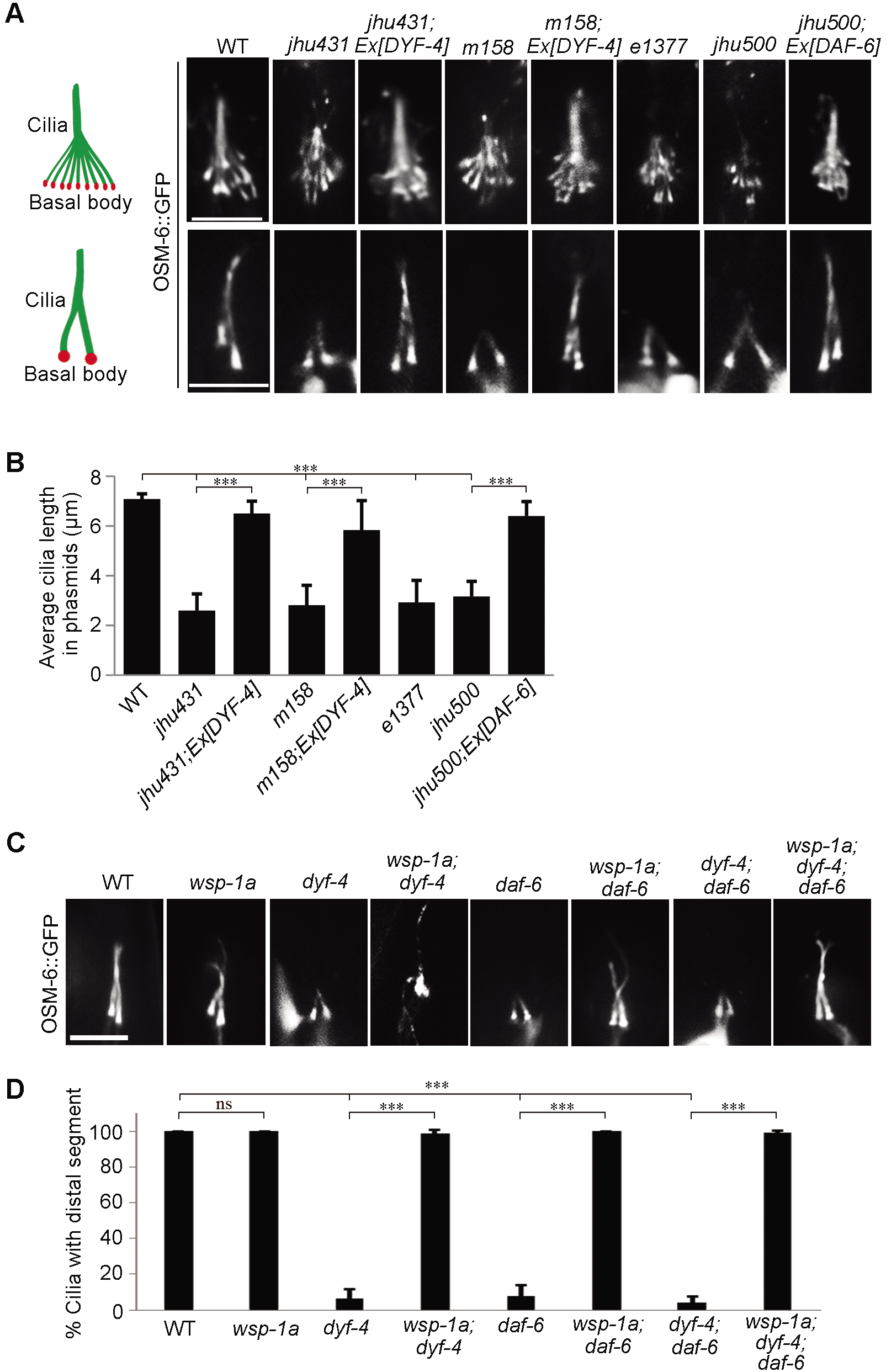
Ciliogenesis is compromised in *dyf-4* and *daf-6* mutants and can be rescued by *wsp-1*. A. Fluorescence micrographs of amphid and phasmid cilia in the indicated worm lines expressing OSM-6::GFP. B. Phasmid cilia length quantification in the indicated worm lines. C. Fluorescence micrographs of phasmid cilia in the indicated genotypes. D. Percentage of severely collapsed dendrites in the phasmid in the indicated worm lines. Scale bars: 5 μm. Data are presented as the mean ± SEM (n ≥ 60 for each genotype). ***P < 0.001 (Student’s t-test).

If the ciliary defects in *dyf-4* or *daf-6* mutants are non-cell autonomous consequences caused by dysfunctional glial cells, glial suppressors of *dyf-4* and *daf-6* might also rescue ciliogenesis defects. Consistent with this hypothesis, deletion of *wsp-1*, the suppressor of glial compartment formation of *daf-6* mutants, fully restored cilia length in *dyf-4* or *daf-6* mutants (Fig. 5C and D).

## Discussion

In this report, through a forward genetic screen, we cloned a new gene, *dyf-4*, that encodes a secreted extracellular protein. The *dyf-4* mutant completely recapitulates the phenotype as *daf-6*, showing abnormal morphogenesis of the glial compartment and truncated dendrites and cilia. DAF-6 is a hedgehog patched-related transmembrane protein that functions on the surface of glial compartment channels to control glial compartment formation (Perens and Shaham, 2005). We further discovered that DYF-4 interacts and colocalizes with DAF-6 along the surface of glial cells in the sensory compartment, and the membrane deposition of DYF-4 and DAF-6 shows interdependence. We thus reveal a novel paradigm that a hedgehog patched-like receptor forms a complex with an extracellular component, and this interaction is critical for the formation of a functional glial compartment.

In *C. elegans*, the proper formation of the sensory compartment comprises the spatiotemporal regulation of glial compartment formation, dendritic tip anchoring and ciliogenesis. Among these stereotyped processes, the formation of a functional glial compartment is indispensable for neuronal dendrite extension and ciliogenesis. It has been reported that the ablation of the sheath glial cells in amphids leads to abnormal dendrite elongation (Singhal and Shaham, 2017). The alteration of the actin cytoskeleton of glial cells regulates cilia biogenesis in a non-cell autonomous manner (Zhu et al., 2017). It is reasonable to speculate that the defects in dendrite extension and ciliogenesis observed in *dyf-4* and *daf-6* mutants are indirectly caused by the abnormal formation of the glial compartment. This notion was supported by our observation that reported glial suppressors of *daf-6* could also suppress the neuronal defects of *dyf-4* and *daf-6* mutants. The extracellular matrix of the sensory compartment is the site of dendrite attachment, where the glia-secreted tectorin-related protein DEX-1 and the sensory neuron-secreted tectorin-related protein DYF-7 form a complex to anchor dendrites (Heiman and Shaham, 2009). Although DYF-4 and DAF-6 are dispensable for DYF-7 localization (our unpublished data), it is very likely that DYF-4 and DAF-6 coordinate the crosstalk between the extracellular matrix composition and the intracellular skeleton network to maintain the proper attachment force generated by the DEX-1-DYF-7 complex.

DAF-6 is mislocalized in *daf-19* mutants, which lack a master transcription factor for ciliogenesis and thus show total loss of cilia (Oikonomou et al., 2011; Perens and Shaham, 2005), indicating that cilia reciprocally regulate the glial compartment formation. It is likely that primary cilia could secrete signals to the lumen of the sensory compartment and impact the morphogenesis and function of glial cells. A plausible model is that DAF-6, the patched-related protein, functions as the receptor on the glial side to receive “signals” and that a hedgehog-related molecule is located in the extracellular matrix and acts as the “signal”. However, despite the presence of many hedgehog-related molecules in *C. elegans* (Aspock et al., 1999), no such hedgehog-related “signal” protein has been identified so far. Although bioinformatics analysis indicates that DYF-4 is not a hedgehog-related protein, our finding that DYF-4 is a glia-secreted protein which interacts with DAF-6 and functions in the same pathway as DAF-6 leads us to propose that DYF-4 may be at least related to the extracellular matrix “signal”. Under normal conditions, DYF-4 recruits and/or stabilizes DAF-6 to control glial compartment formation. If external matrix conditions are abnormal, for example, due to abnormal dendrites or cilia, the interaction between DYF-4 and DAF-6 may be compromised, leading to abnormal glial compartment formation. Further detailed investigations of the biochemical characteristics of DYF-4 are needed to determine whether DYF-4 is the ligand of DAF-6 and how DYF-4 regulates DAF-6. Whether DYF-4 is the ligand of DAF-6 or not, studies on DYF-4 and its interacting proteins will certainly provide new avenues for understanding the formation of sensory compartments.

## Materials and Methods

### *C. elegans* strains

The *C. elegans* strains used in this study are listed in Key Resources Table 1. The worms were cultured and maintained on NGM agar seeded with OP50 bacteria at 20°C under standard conditions. The *dyf-4(jhu431)* and *daf-6(jhu500)* mutants were identified from an EMS screening for mutants showing dye filling defects. Standard genetic crossing was used to introduce reporter transgenes into worms of various genetic backgrounds and generate double or triple mutants. Genotyping was performed using PCR.

### Dye-filling assay

Worms were rinsed into a 1.5 ml EP tube with M9 buffer, followed by centrifugation in a desktop centrifuge at low speed. Then, the supernatant was removed, and the worms were washed twice with M9 buffer. After washing, the worms were incubated with 10 μg/ml DiI in M9 buffer at room temperature for 2 h and were then washed again with M9 buffer three times. Dye filling in the amphid was observed with a Nikon SMZ18 stereomicroscope, and dye filling in the phasmid was observed with a Nikon Eclipse Ti microscope with a Plan Apochromat 100× objective.

### Plasmids and transgene generation

To generate the *Pdyf-4::DYF-4::GFP* plasmid, a DNA fragment including the 1511 bp promoter and the whole genomic DNA sequence of *dyf-4* was cloned into the SmaI and KpnI restriction sites of the GFP vector pPD95.75. For the construction of *Pdyf-4::DYF-4^C366Y^::GFP*, the guanine (G) nucleotide at site 1097 of the *dyf-4* CDS was replaced by adenosine (A). For the construction of *Pdyf-4::GFP*, 1511 bp of the promoter of *dyf-4* was amplified by PCR and ligated into the SmaI and KpnI restriction sites of pPD95.75. To generate *Pdaf-6::DAF-6::GFP*, 3096 bp of the promoter of *daf-6* was ligated into the BamHI and SmaI restriction sites of pPD95.75, and the CDS of *daf-6* was ligated into the SmaI and KpnI restriction sites of pPD95.75. To construct *Pdaf-6::DYF-4::GFP* and *Pdyf-7::DYF-4::GFP*, the *dyf-4* CDS was ligated into the SmaI and PstI restriction sites of pPD95.75, and 3096 bp of the promoter of *daf-6* or 1758 bp of the promoter of *dyf-7* was ligated into the PstI and SmaI restriction sites of pPD95.75.

### Microscopy and imaging

Worms were anaesthetized using 20 mM levamisole, mounted on 4% agar pads and then imaged by using a Nikon Eclipse Ti microscope or Olympus FV1000a confocal microscope. L4 worms were used to record phasmid dendrite length statistics. The fluorescence intensity was measured using NIS-Elements software. The relative fluorescence intensity of cilia was calculated by subtracting the average fluorescence intensity outside the cilia from the average fluorescence intensity inside the cilia.

### Immunofluorescence of DYF-4::GFP

Worms harboring the DYF-4::GFP construct were placed on a glass slide and squashed with a coverslip. Then, the slide was frozen in liquid nitrogen, and the coverslip was removed. Then, the sample was fixed in −20°C methanol for 20 min, followed by fixation in −20°C acetone for 10 min, washing in PBS, blocking in 3% BSA, and sequential incubation with an anti-GFP primary antibody (mouse anti-GFP, 1:200, 11814460001, Roche) and a corresponding secondary antibody (goat anti-mouse Alexa Fluor 488). A Nikon Eclipse Ti-E microscope was used for observation and imaging.

### GST pull-down assay

Plasmids for the expression of His-tagged DAF-6 fragments or GST-tagged DYF-4 were constructed using pET28a or pGEX-4T-1, respectively. All plasmids were transfected into *Escherichia coli* strain BL21 (DE3) to express the proteins. Protein expression was induced with 0.5 mM IPTG at 18°C overnight, and the proteins were then purified by using Ni-Sepharose beads (GE Healthcare) or GST Sepharose beads (GE Healthcare). GST pull-down was performed as described previously (Wei et al., 2013). Briefly, the purified His-DAF-6 ECL1 or His-DAF-6 ECL4 protein was incubated with GST-DYF-4 or GST immobilized on glutathione Sepharose beads in binding buffer (50 mM Tris-HCl, pH 7.4, 150 mM NaCl, 1% Triton X-100, 1 mM dithiothreitol, 10% glycerol, protease inhibitors) for 4 h at 4°C. After incubation, beads were washed 5 times with the binding buffer, after which the loading buffer was added, and the sample was boiled for 10 min. The samples were then subjected to SDS-PAGE and analyzed by western blotting with a monoclonal antibody against His.

### Statistical analysis

Statistical differences between two samples were determined using the two-tailed unpaired Student’s *t*-test in Excel or GraphPad Prism 5 software. P values >0.05 were considered to indicate a nonsignificant difference. P values < 0.001 (marked as ***) were considered to indicate a significant difference.

## Acknowledgements

We thank the *Caenorhabditis Genetics* Center, the Japanese National Bioresource Project, and Dr. Guangshuo Ou for worm strains. This work was supported by National Natural Science Foundation Youth Project of China (Grant 31702019) to H.H., and National Natural Science Foundation of China (Grant 31671549) to Q.W.

## Author Contributions

Under the supervision of Q. W., H. H. and H. C. performed most of the experiments, collected and analyzed the data. *jhu431* and *jhu500* were isolated in J. H. and K. L. lab. Y. Z. and Q. W. cloned *jhu431* mutant. Z. W., Y. Z., Z. H. and J. Z. assisted the experiments. Q.W. and J. H. wrote the paper with the help of H.H., H.C. and K. L.

## Supplementary Figure legends

**Figure 1-figure supplement 1.**
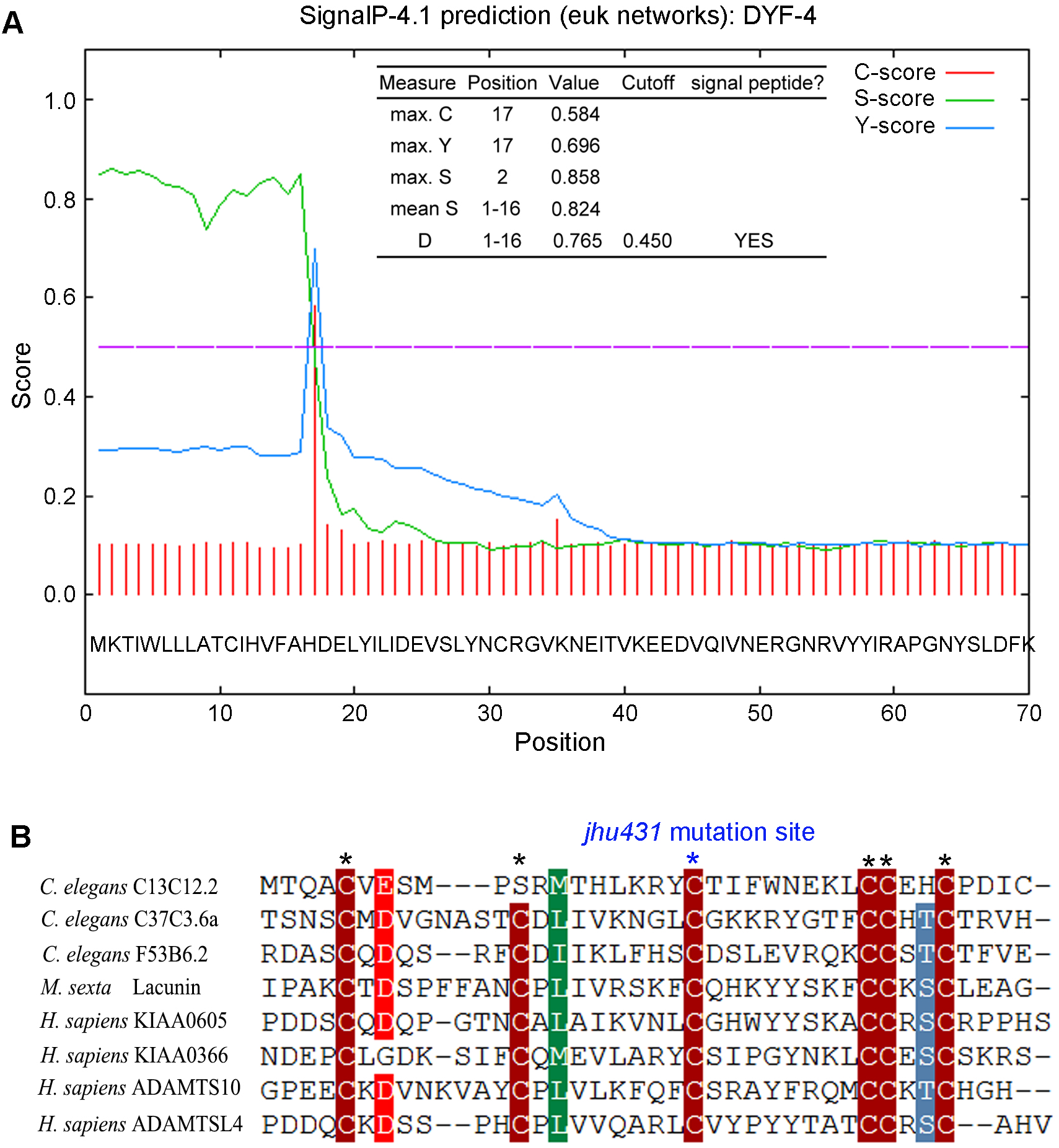
Protein structure prediction for DYF-4. A. Signal peptide prediction for DYF-4 by SignalP4.1. B. Sequence alignment of the PLAC domain. *, the conserved cysteines in the PLAC domain.

**Figure 2-figure supplement 1.**
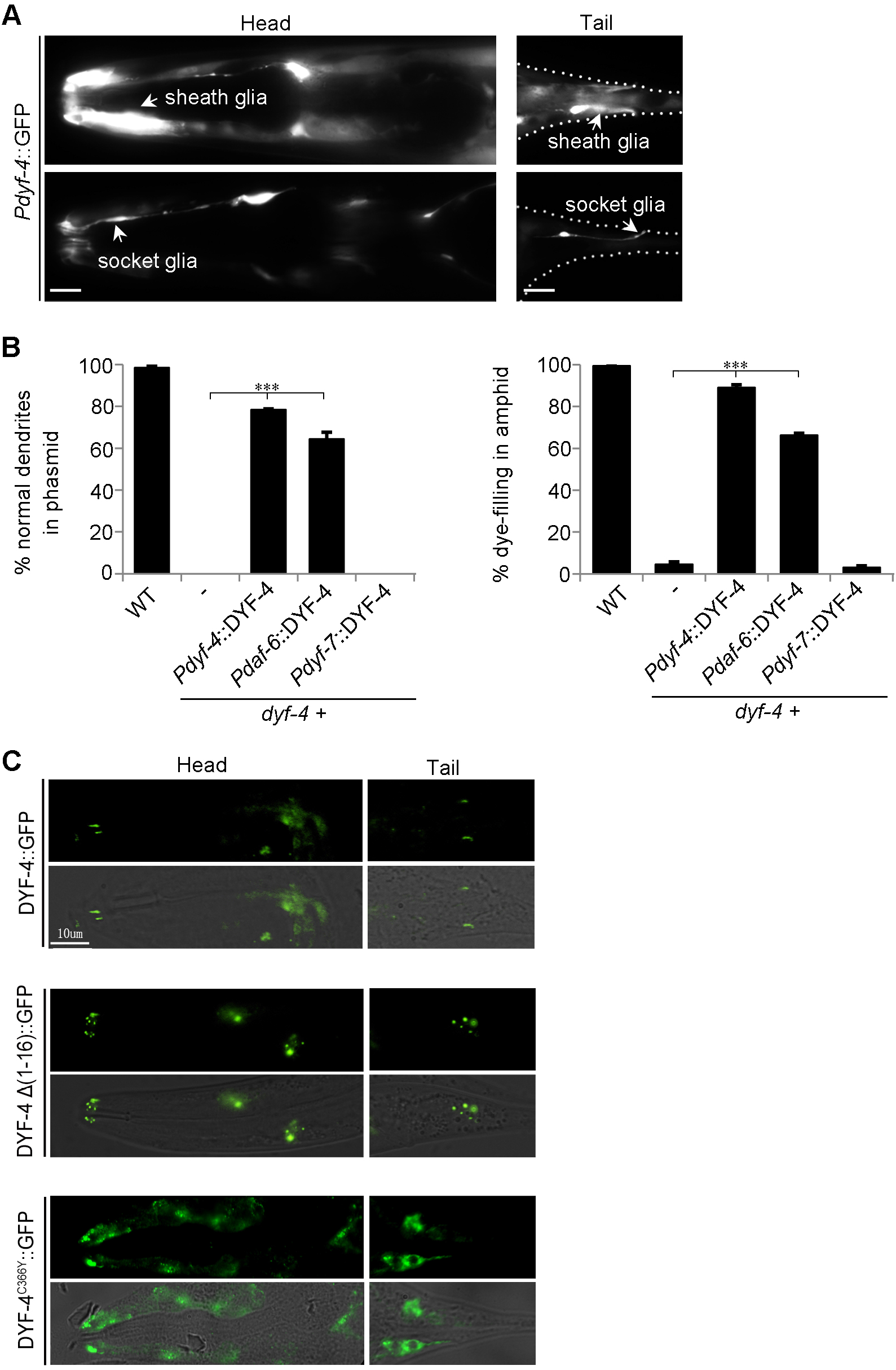
DYF-4 is expressed in glial cells. A. Fluorescence micrographs of the *dyf-4* expression pattern in the head and tail region in worms expressing *Pdyf-4*::GFP. Sheath glial cells and socket glial cells are indicated by arrowheads. Scale bar: 5 μm. B. The percentages of normal dendrites in the phasmid (left) and dye filling in the amphid (right) in WT and *dyf-4(m158)* mutants expressing the WT *dyf-4* gene driven by different promoters. Data are presented as the mean ± SEM from three independent experiments (n ≥ 80 per experiment). ***P < 0.001 (Student’s t-test). C. The subcellular localization pattern of DYF-4 (C366Y) and DYF-4 Δ (1-16). Scale bar: 10 μm.

**Figure 3-figure supplement 1.**
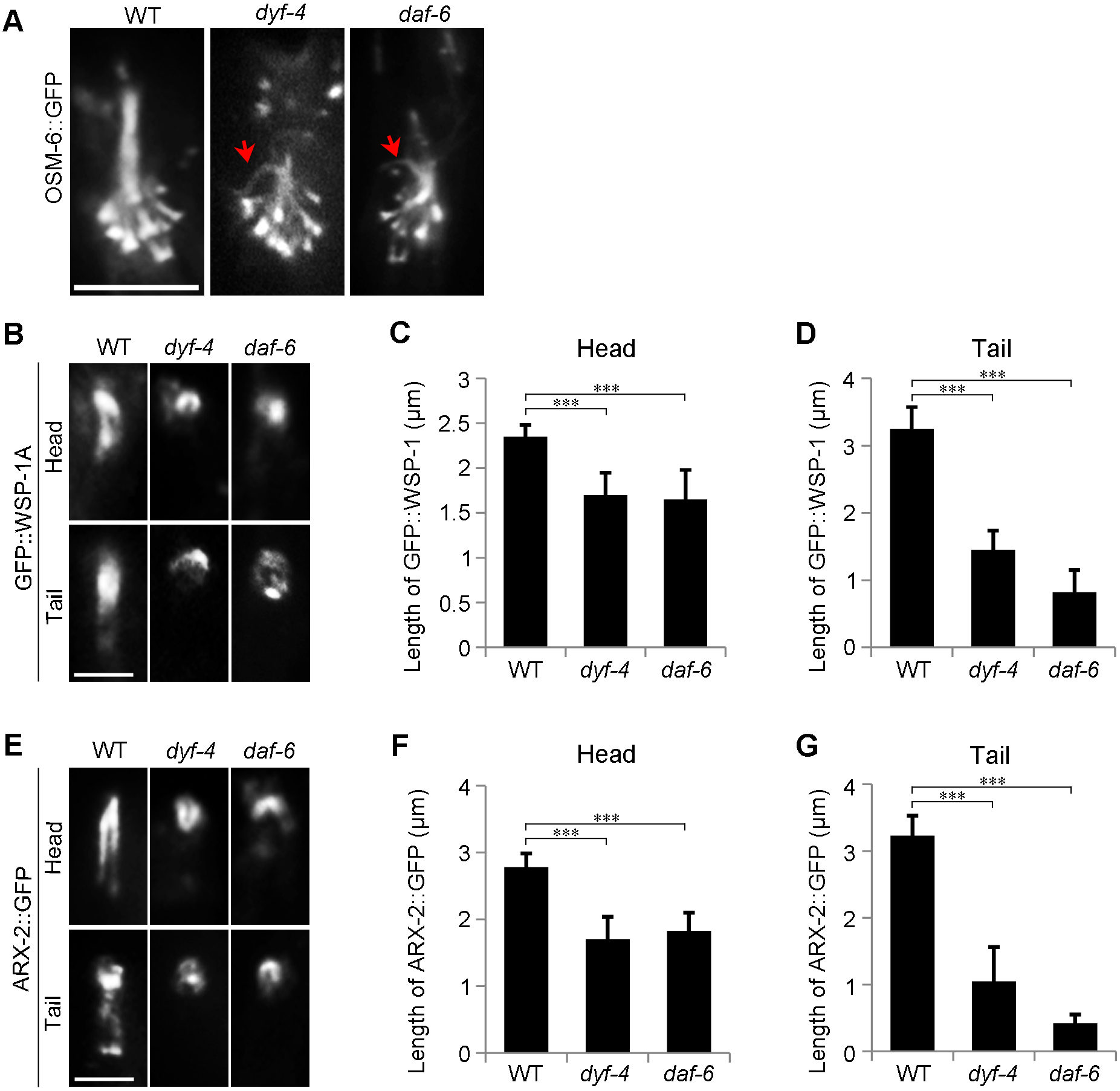
The glial compartment is compromised in *dyf-4* mutants. A. Fluorescence micrographs of curved cilia in *dyf-4* and *daf-6* mutant worms. Curved cilia are indicated by red arrowheads. Scale bars = 5 μm. B. WSP-1A localization at the head and tail in WT, *dyf-4(m158)* and *daf-6(e1377)* worms. Scale bars: 2 μm. C. Quantification of GFP::WSP-1A signal length in the amphids of WT, *dyf-4(m158)* and *daf-6(e1377)* worms. D. Quantification of GFP::WSP-1A signal length in the phasmids of WT, *dyf-4(m158)* and *daf-6(e1377)* worms. E. ARX-2 localization at the head and tail in WT, *dyf-4* and *daf-6(e1377)* worms. Scale bars: 2 μm. F. Quantification of ARX-2::GFP signal length in the amphids of WT, *dyf-4(m158)* and *daf-6(e1377)* worms. G. Quantification of ARX-2::GFP signal length in the phasmids of WT, *dyf-4(m158)* and *daf-6(e1377)* worms. All data are presented as the mean ± SEM (n ≥ 50 for each genotype). ***P < 0.001 (Student’s t-test).

**Figure 5-figure supplement 1.**
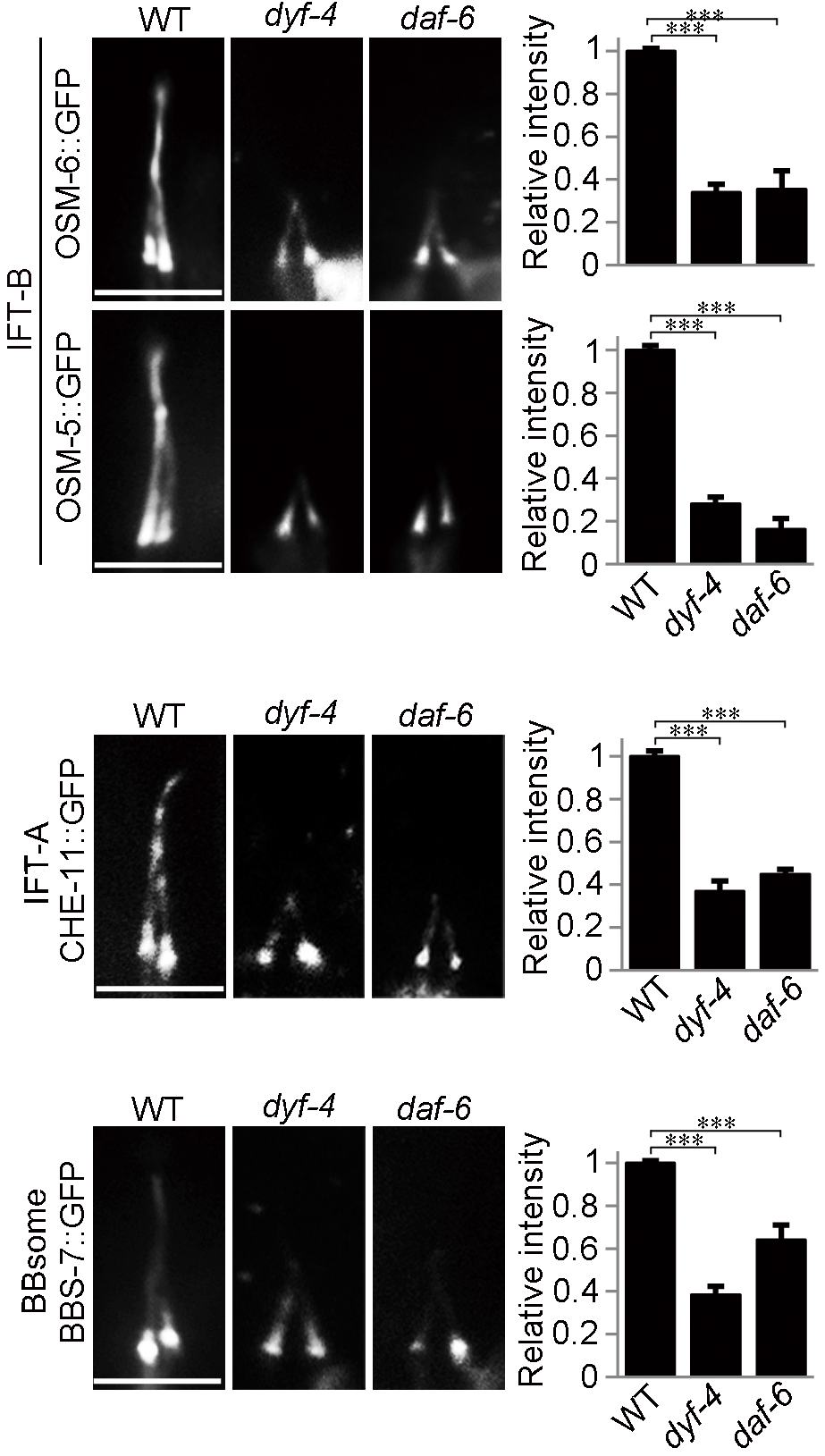
The localization of IFT components in WT, *dyf-4* and *daf-6* worms. Fluorescent micrographs and quantification of the relative fluorescence intensities in the phasmid cilia of WT, *dyf-4(m158)* and *daf-6(e1377)* worms expressing various IFT markers: IFT-B components OSM-6::GFP and OSM-5::GFP, IFT-A component CHE-11::GFP and BBsome component BBS-7::GFP. Data are presented as the mean ± SEM (n ≥ 60 for each genotype). ***P≤0.001 (Student’s t-test). Scale bars: 5 μm.

**Key Resources Table 1. Worm strains used in this study.**

